# Targeting wild-type IDH1 enhances chemosensitivity in pancreatic cancer

**DOI:** 10.1101/2023.03.29.534596

**Authors:** Mehrdad Zarei, Omid Hajihassani, Jonathan J. Hue, Hallie J. Graor, Luke D. Rothermel, Jordan M. Winter

**Affiliations:** Case Comprehensive Cancer Center, Case Western Reserve University, Cleveland, OH; Department of Surgery, Division of Surgical Oncology, University Hospitals Cleveland Medical Center, Cleveland, OH

**Author notes:** Corresponding Author: Jordan M. Winter, MD University Hospitals Cleveland Medical Center, Department of Surgery 11100 Euclid Ave. Cleveland, OH 44106.

**Keywords:** Pancreatic cancer, IDH1, ivosidenib, targeted therapy chemotherapy, chemoresistance, combination therapy

## Abstract

Pancreatic cancer (PC) is one of the most aggressive types of cancer, with a five-year overall survival rate of 11% among all-comers. Current systemic therapeutic options are limited to cytotoxic chemotherapies which have limited clinical efficacy and are often associated with development of drug resistance. Analysis of The Cancer Genome Atlas showed that wild-type isocitrate dehydrogenase (wtIDH1) is overexpressed in pancreatic tumors. In this study, we focus on the potential roles of wtIDH1 in pancreatic cancer chemoresistance. We found that treatment of pancreatic cancer cells with chemotherapy induced expression of wtIDH1, and this serves as a key resistance factor. The enzyme is protective to cancer cells under chemotherapy-induced oxidative stress by producing NADPH and alpha-ketoglutarate to maintain redox balance and mitochondrial function. An FDA-approved mutant IDH1 inhibitor, ivosidenib (AG-120), is actually a potent wtDH1 inhibitor under a nutrient-deprived microenvironment, reflective of the pancreatic cancer microenvironment. Suppression of wtIDH1 impairs redox balance, results in increased ROS levels, and enhances chemotherapy induced apoptosis in pancreatic cancer vis ROS damage *in vitro*. *In vivo* experiments further revealed that inhibiting wtIDH1 enhances chemotherapy anti-tumor effects in patient-derived xenografts and murine models of pancreatic cancer. Pharmacologic wtIDH1 inhibition with ivosidenib represents an attractive option for combination therapies with cytotoxic chemotherapy for patients with pancreatic cancer. Based on these data, we have initiated phase Ib trial combining ivosidenib and multi-agent chemotherapy in patients with pancreatic cancer (<u>NCT05209074</u>).

## Introduction

While surgical resection plays a key role in the management of patients with localized pancreatic cancer (PC), a fraction of all patients present with surgically resectable disease (1). In fact, PC is often considered a systemic disease at the time of diagnosis, even without radiographic evidence (2). Thus, systemic therapies are arguably the most important treatment modality for all comers with PC, with hopes of controlling micro- and macro-metastatic disease. Current systemic therapeutic options are limited to cytotoxic chemotherapies with minimal clinical efficacy. FOLFIRINOX a combination of chemotherapy drugs (folinic acid, 5-fluorouracil, irinotecan, oxaliplatin), was initially described in 2011 (3,4), and is still considered first-line therapy over a decade later with best clinical benefit in pancreatic cancer patients(5,6). Yet, this regimen only improves median survival by approximately 4 months for patients with metastatic disease relative to single-agent gemcitabine (7–10). It is a harsh reality that most patients with metastatic PC will not survive one year. Identification of novel, effective therapeutics represents a critical area of research.

Recent investigations by our lab have identified the metabolic enzyme wild-type isocitrate dehydrogenase (wtIDH1) as a key enzyme that drives important pro-survival mechanisms in PC. IDH1 is a cytosolic enzyme that is isofunctional to the mitochondrial enzymes IDH2 and 3A. The enzyme catalyzes the interconversion of isocitrate and alpha-ketoglutarate (αKG) using NADP(H) as a cofactor (11–13). Under nutrient limitations commonly present in the tumor microenvironment, oxidative decarboxylation of isocitrate is favored, producing NADPH and αKG. These products directly support an antioxidant defense and mitochondrial function (through anaplerosis into the TCA cycle), respectively (14–16). Our recent studies in PC identified pharmacologic inhibition of wtIDH1 dramatically improves survival in multiple murine models of PC (14). Ivosidenib (AG-120) is approved by the United States Food and Drug Administration for the treatment of mutant IDH1 cholangiocarcinoma and select hematologic malignancies (17–19); however, our work validated AG-120 and other small molecules inhibitors of mutant IDH1 are actually potent wild-type IDH1 inhibitor (20–22) in conditions encountered in the tumor microenvironment (e.g., low nutrient levels). Importantly, ivosidenib is well tolerated and administered orally, making it an attractive option for combination regimens. IDH1 plays a key role in redox homeostasis. As chemotherapies are known inducers of reactive oxygen species (23), pharmacologic inhibition of IDH1 may prove to be a synthetic lethal combination when coupled with cytotoxic agents.

Herein, we show that wtIDH1 is especially important for PC cell survival, particularly when exposed to chemotherapy. Further, we trial ivosidenib combined with common chemotherapies in murine models of pancreatic cancer to identify optimal multi-agent regimens for use in future clinical trials.

## Materials and Methods

### Cell lines and cell culture

Human pancreatic cancer cell lines, MiaPaca2 and PANC-1, were obtained from ATCC (American Type Culture Collection) (Rockville, MD, USA). Murine pancreatic cell, KPC (K8484: KrasG12D; Trp53R172H/+; Pdx1-Cre), were gifts from Dr. Darren Carpizo. All cell lines were cultured in normal DMEM with 10% fetal bovine serum (Gibco/Invitrogen) and 1% penicillin/streptomycin (Invitrogen). Cell lines were maintained at 37°C in 5% humidified CO_2_. Glucose-free DMEM (Life Technologies, 21013-024) supplemented with varying glucose concertations was used for low glucose experiments, and magnesium sulfate-depleted DMEM (Cell Culture Technologies, 964DME-0619) was used for low magnesium experiments. The low glucose and low magnesium media were typically used to simulate nutrient withdrawal. All cell lines were Mycoplasma-tested routinely using a mycoplasma detection kit (ATCC, 30-1012K). Before any experiment, cells were passaged at least twice.

### Immunohistochemistry (IHC)

The tumors were fixed overnight in 10% formalin and then switched to PBS. IHC was performed on 5 µm-thick formalin-fixed paraffin-embedded (FFPE) tissue sections on glass slides. Tissue sections were incubated for 75 minutes at 60 °C, followed by a series of washes for deparaffinization and then rehydrated. Antigen retrieval was performed using a pressure cooker for 10 minutes at 123 °C in 10 mM citrate at pH 6.0, then 3% H_2_O_2_ for 10 minutes, followed by washing with water. The tissue sections were then incubated with a blocking solution containing 5% goat serum (Cell Signaling Technology; 5425) at room temperature for 1 hour. The primary antibodies were diluted (SignalStain Antibody Diluent; 8112) and incubated overnight at 4 °C using the following antibodies: Ki67 (SP6, BioCare Medical) and cleaved caspase 3 (Cell Signaling Technology; 9579). The slides were washed thrice by 1X TBST and incubated using IHC detection Signal Stain Boost IHC Detection Reagent (Cell Signaling Technology; 8114) at room temperature for 30 minutes. The slides were washed three times with 1X TBST. Signal was detected by the HRP substrate kit (Vector; SK-4200), according to the manufacture’s recommendations. Slides were counterstained with hematoxylin before final dehydration, and coverslips were mounted. Quantification of each staining was performed by using ImageJ software at 20x magnification.

### Cell growth assays

Cells were seeded in 96-well plates at 2,000 cells per well, and after 24 hours, cells were treated with chemotherapies including 5FU (Sigma, F6627), oxaliplatin (Sigma, O9512), and IDH1 inhibitor AG-120 (Asta Tech, 40817) at indicated concentrations. All experiments for cell viability lasted for 6 days unless otherwise detailed. Cell viability was measured using the Quant-iT^TM^ PicoGreen^TM^ dsDNA assay kits (Invitrogen; P7589).

Drug combination assays were performed by seeding 1,000-2,000 cells per well in 96-well plates. After 24 hours, cells were treated with different concentrations of AG-120 (dose range, 1.25-20 μM/mL), 5-FU (dose range, 0.15-20 μM/mL) in a 6 x 8 matrix. Each treatment was done in triplicate. Cells were treated for 6 days, and the cell growth relative to vehicle treatments was measured by staining with Quant-iT PicoGreen. For all *in vitro* experiments using AG-120, cells were cultured under low magnesium conditions (<0.4 mM Mg^2+^) to effectively inhibit wtIDH1 enzyme activity (as a reference, normal culture media and serum contain roughly 1 mM Mg^2+^). Low glucose (2.5 mM glucose) was utilized as indicated to generate conditions of wtIDH1 dependency and simulate glucose levels in the tumor microenvironment (24–27).

Drug interaction data were quantified and characterized as synergistic, additive, or antagonistic, as described previously (28).

### Immunoblot analysis

Cells were lysed using 1X ice-cold RIPA buffer supplemented with protease and phosphatase inhibitors. Protein concentration was quantified using the Pierce™ B.C.A. Protein Assay kit (Thermo Fischer Scientific). Equal amounts of whole protein extracts were loaded and separated by electrophoresis on a 4–12% Bis-Tris gel (Life Technologies) and transferred to a PVDF membrane (Thermo Fischer Scientific). Blots were blocked in 5% (wt/vol) non-fat milk and then probed with antibodies against anti-*IDH1* (Invitrogen, OTI2H9) and anti-beta-actin (Invitrogen, 15739-BTIN). Chemiluminescent (32106, Thermo Fisher Scientific) signal was captured using a digital imager (Odyssey Infrared Imaging system).

### Clonogenic assay

Cells were cultured and seeded in 6-well plates at 3,000-5,000 per well depending on the growth rate in 2 mL media to assess the clonogenicity of cells. The media was not changed during experiments unless indicated. Cells were treated with AG-120 or 5-FU after being cultured with low glucose level media followed by low MgSO4 (0.08 mM). After 8-10 days upon completing the experiments, cells were fixed in a reagent containing 80% methanol and stained with 0.5% (w/v) crystal violet solution for 10 minutes (29). To determine the relative proliferation of the cells, the dye was extracted using 10% glacial acetic acid, and the associate absorbance was measured at 600 nm by a Promega GloMax plate reader.

### Cellular ROS and apoptosis analysis

Cells were plated in 96-well black plates. Cellular reactive oxygen species (ROS) levels were detected using 2’,7’ dichlorodihydrofluorescein diacetate (H2DCFDA; Invitrogen). Cells were treated in the absence or presence of drugs for 48 hours. Then incubated with 100 μL phenol red-free media containing 10 μM DCFDA for 45 minutes at 37 °C in the dark. The fluorescence signal was detected using an excitation wavelength at 485 nm and emission wavelength at 535 nm on a GloMax Promega plate reader. For apoptosis, cells were plated in white 96-well plates and treated for 48 hours in the presence of drugs. The caspase 3/7 (Caspase-GloTM Promega G8090) level was measured per the manufacturer’s instructions.

### Real time quantitative PCR

Total RNA was extracted using the RNeasy PureLink RNA isolation (Life Technologies; 12183025) according to manufacturers’ instructions. RNA was converted to cDNA using a High-Capacity cDNA Reverse Transcription Kit following the manufacturer’s protocol (Applied Biosystems; 4387406). The qPCR was performed using Taqman™ Universal Master Mix (Thermo Fisher Scientific; 4440038) with probe (IDH1 Thermo Fisher Scientific; 4351372) and cDNA and was analyzed using the Bio-Rad CFX96 Maestro manager 2.0 software.

### Bioenergetics analysis

MiaPaca2 and PANC-1 cells were seeded at 10,000 cells/well in complete normal DMEM (25 mM Glucose and 2 mM Glutamine) in an Agilent XFp Cell Culture miniplate (#103025-100) at 37°C in 5% humidified CO_2_ incubators, as recommended. OCR was performed using the XFp mini extracellular analyzer (Seahorse Bioscience). For OCR, cells were treated in the absence or presence of 5-FU under indicated glucose concentration for 36 hours. The XFp FluxPak cartridge (#103022-100), was hydrated with 200 μL/well of XF Calibrant (#100840-000) and incubated at 37□°C using non-CO_2_ incubator overnight. The following day cells were washed and replaced with Seahorse XF base media (at indicated glucose concentrations), and the plate was incubated in a non-CO2 incubator at 37°C. The Mito Stress test kit (#103015-100) was reconstituted in an assay medium to make the following concentration of inhibitors: 1.5 μM oligomycin, 2.5 μM FCCP, and 0.5 μM rotenone□+□0.5 μM antimycin A. OCR was measured using Wave 2.6 software, according to the manufacturer’s instructions, all data was normalized to cell number with Quant-iT PicoGreen^TM^ dsDNA Assay Kit (Invitrogen).

### Animal studies

All mice experiments were approved and conducted under the Case Western Reserve University (CWRU) Institutional Animal Care Regulations and Use Committee (IACUC, protocol 2018-0063) guidelines. Female athymic nude mice (Nude-Foxn1nu) (aged six to eight weeks) were purchased from Harlan Laboratories (6903M). Patient-derived xenografts (PDX) were purchased from Jackson Laboratory (#TM01212) and 2 mm^3^ tumors were implanted subcutaneously in the right flanks of nude mice. Mia-Paca2 cells were suspended in 200 μL solution comprised of 70% Dulbecco’s PBS and 30% Matrigel. Suspension of 1×10^6^ cells was injected subcutaneously into the right flank of mice. Ten-week-old C57BL/6J female mice were purchased from Jackson Laboratories for orthotopic experiments. A suspension of 5×10^4^ Luciferase-expressing KPC cells in 30 μL of a 40:60 mixture of PBS and Matrigel solution (Corning, 354234) was injected carefully into the pancreas. On postoperative day 10, the presence of pancreatic tumors was confirmed by bioluminescence (BLI) via Spectrum CT (PerkinElmer, 2898979) following an intraperitoneal injection of 100 μL D-luciferin (50 mg/mL in PBS). Mice with confirmed tumors were randomized to indicated treatment groups. When the PDX and Mia-Paca2 tumor volumes reached 100-120 mm^3^ (approximately 3-5 weeks) or after orthotopic tumors were confirmed with BLI, treatments were initiated to the indicated conditions. Mice were administered AG-120 (Asta Tech, 40817; 75 mg/kg in PEG-400, Tween-80, and saline (10:4:86) twice a day orally), 5-Fluorouracil (Sigma-Aldrich, F6627; 30 mg/kg in sterile saline via intraperitoneal injections two times a week), oxaliplatin (Sigma-Aldrich, O9512; 5 mg/kg), FOLFIRINOX (5FU 12.5 mg/kg, irinotecan 25 mg/kg (Sigma-Aldrich, I1406) and oxaliplatin 2.5 mg/kg, once weekly). or vehicle control.

Body weight and tumor volume were measured weekly. Vernier calipers were used to measure tumor size, using the formula (Volume = Length × Width^2^/2). Upon termination of the experiment, the mice were euthanized under carbon dioxide inhalation and tumors were resected. The tumors were collected in 10% formalin (Fisher brand; 427-098) for IHC and stored at -80°C until processed for further analysis.

### Statistical analysis

Data were expressed as mean ± SEM (standard error of the mean) of at least three independent experiments. Comparisons between groups were determined using unpaired, two-tailed Student *t-*test (* *p* < 0.05; ** *p* < 0.01; *** *p* < 0.001 **** *p* < 0.0001). One-way or Two-way ANOVA test was used to compare more than two groups. The Kaplan-Meier estimate was used for survival experiments, and groups were compared using the log-rank test. GraphPad Prism 9.2.3 Software was used for statistical analysis.

## Results

### Wild-type IDH1 is overexpressed in PC and promotes chemotherapy resistance

The Cancer Genome Atlas (TCGA) database analysis revealed that wtIDH1 is highly expressed in tumor samples from patients with PC. Further, higher tumoral expression of IDH1 was associated with poor survival, relative to lower expression, among patients with PC in TCGA **(Fig. 1A and B)**. The development of chemoresistance in PC severely limits the effectiveness of chemotherapeutics. Herein, to investigate a possible relationship between IDH1 expression and chemotherapeutic drugs, PC cells were treated with 5-FU. Using qPCR and immunoblot, 5-FU induced expression of IDH1 in both Mia-Paca2 and PANC-1 cells (**Fig. 1C and D**). These data demonstrate that IDH1 expression is strongly associated with chemotherapeutic treatment.

**Figure-1.**
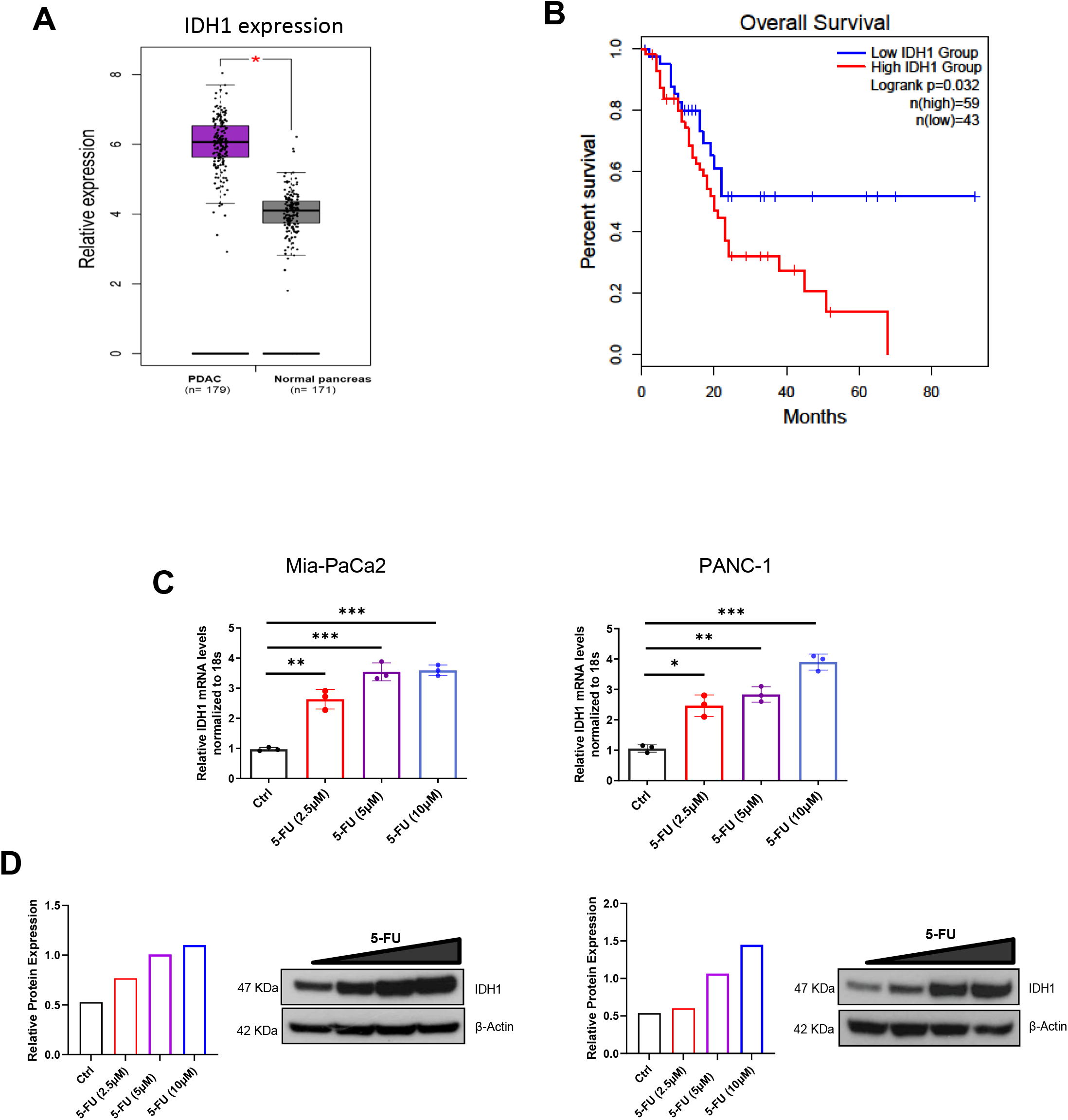
Wild-type IDH1 confers to chemoresistance. **A]** RNA-sequencing data obtained from TCGA showing expression of IDH1 in pancreatic cancer compared to normal pancreatic tissue *P < 0.05. **B]** Association between wtIDH1 expression and overall survival, analyzed using Kaplan-Meier estimate and log-rank test; P=0.032. **C]** qPCR analysis of wtIDH1 in Mia-Paca2 and PANC-1 cells after 48 hours of treatment with different concentrations of 5-FU (2.5, 5, and 10 μM) **D]** Immunoblotting analysis of wtIDH1 expression in Mia-Paca2 and PANC-1 cells after 72 hours of treatment with different concentration of 5-FU (2.5, 5 and 10 μM). Each data point represents the mean ± SEM of three independent experiments. *, P *<* 0.05; **, P *<* 0.01; ***, P *<* 0.001.

### Targeting IDH1 sensitizes PC cells to chemotherapy *in vitro*

5-FU is often used in multi-agent chemotherapy regimens for patients with PC (30), as it was ineffective as a monotherapy in historic trials (31,32). Previously our group validated AG-120 as a potent wtIDH1 inhibitor under low magnesium and low nutrient conditions present in the PC tumor microenvironment (14). As IDH1 mRNA and protein levels increased following chemotherapy treatment, we hypothesized that IDH1 inhibition would synergize with chemotherapies to augment cytotoxic effects. To study the role of IDH1 in chemoresistance, we treated the Mia-Paca2 and PANC-1 cells with AG-120 and chemotherapies. The dose-responses of each drug as a monotherapy **(Supplementary Fig. S1A -S1B)** informed the optimal dosing doses for combination studies. IDH1 inhibition with AG-120 rendered 5-FU substantially more potent (up to 31-fold and 11-fold at some dosing levels) in both Mia-Paca2 and PANC-1 cell lines **(Fig. 2A and C)**. The Bliss index analysis data showed that IDH1 inhibition in combination strongly synergizes with 5-FU, and the combination is mostly synergistic in both Mia-Paca2 and PANC-1 cell lines, as seen in the Bliss matrix **(Fig. 2B and D)**.

**Figure-2.**
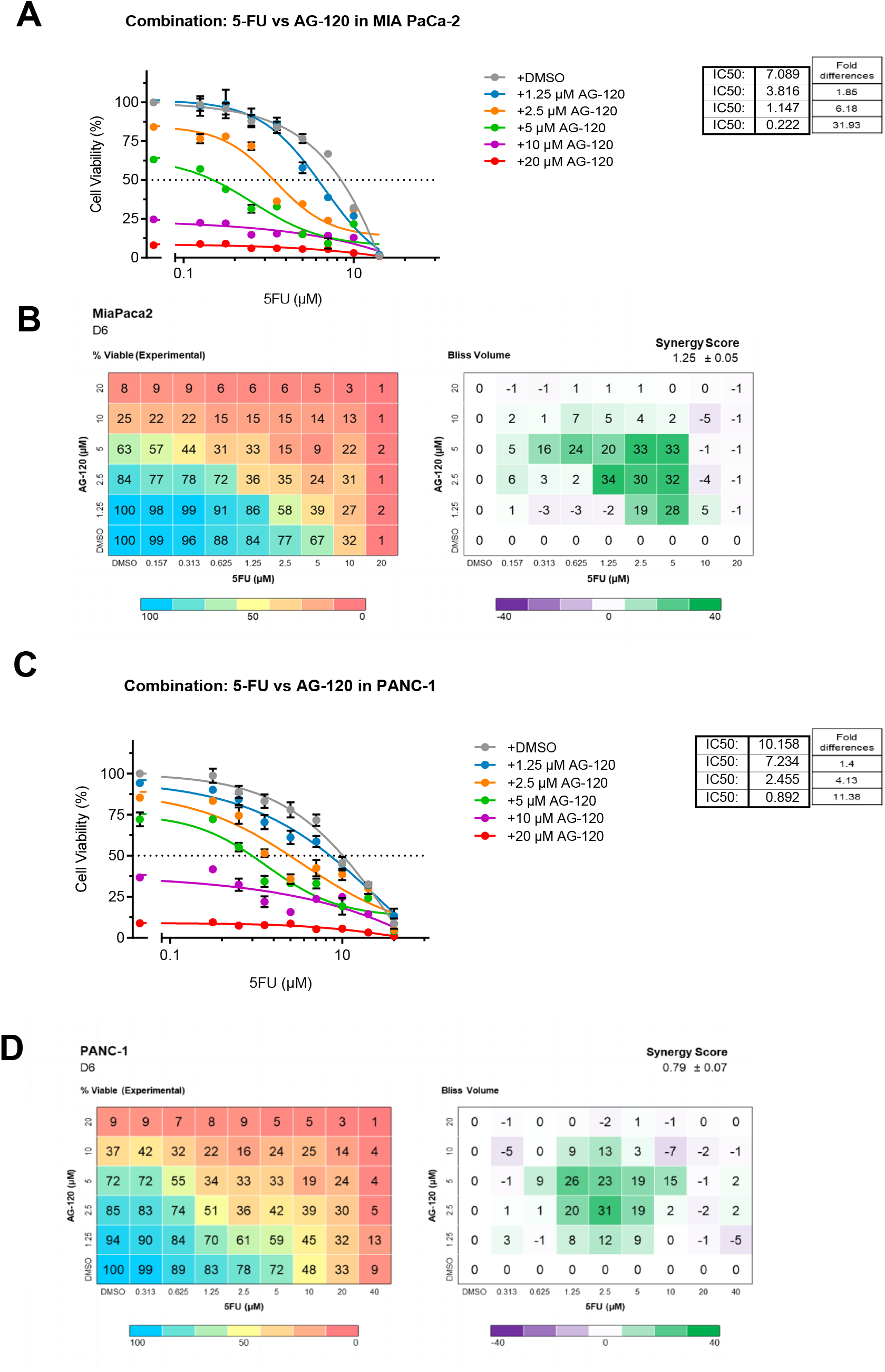
Targeting IDH1 sensitizes PC cells to chemotherapy. **A] and C]** Drug sensitivity in Mia-Paca2 and PANC-1, with varying concentrations of 5-FU and AG-120, cultured for 5-6 days under low glucose. IC_50_ results and fold differences are provided. **B] and D]** Drug matrix heatmap 6 x 8 (AG-120 and 5-FU) grid showing percent viability and Bliss Independence scores in Mia-Paca2 and PANC-1 cells cultured under low glucose for 5-6 days. Positive values reflect synergy and appear green on the heatmap. All treatments with AG-120 were carried out under low glucose and low Mg^2+^ concentrations.

Next, we performed clonogenic assays to understand the long-term effects of IDH1 inhibition in combination with 5-FU on PC cells. Both Mia-Paca2 and PANC-1 cells were treated at increasing doses of drugs, and colonies were stained after 10 days **(Fig. 3)**. Both AG-120 and 5-FU as monotherapy at low doses did not significantly reduce colony formation of PC cells (**Fig. 3A and B**). We further tested the combination of AG-120 and 5-FU in both cell lines. As above, inhibition of IDH1 with AG-120 sensitizes Mia-Paca2 and PANC-1 to 5-FU **(Fig. 3 A and B)**. Together these data indicate that IDH1 inhibition has efficacy against PC cells and can be utilized to increase the sensitivity of 5-FU for the treatment of PC.

**Figure-3.**
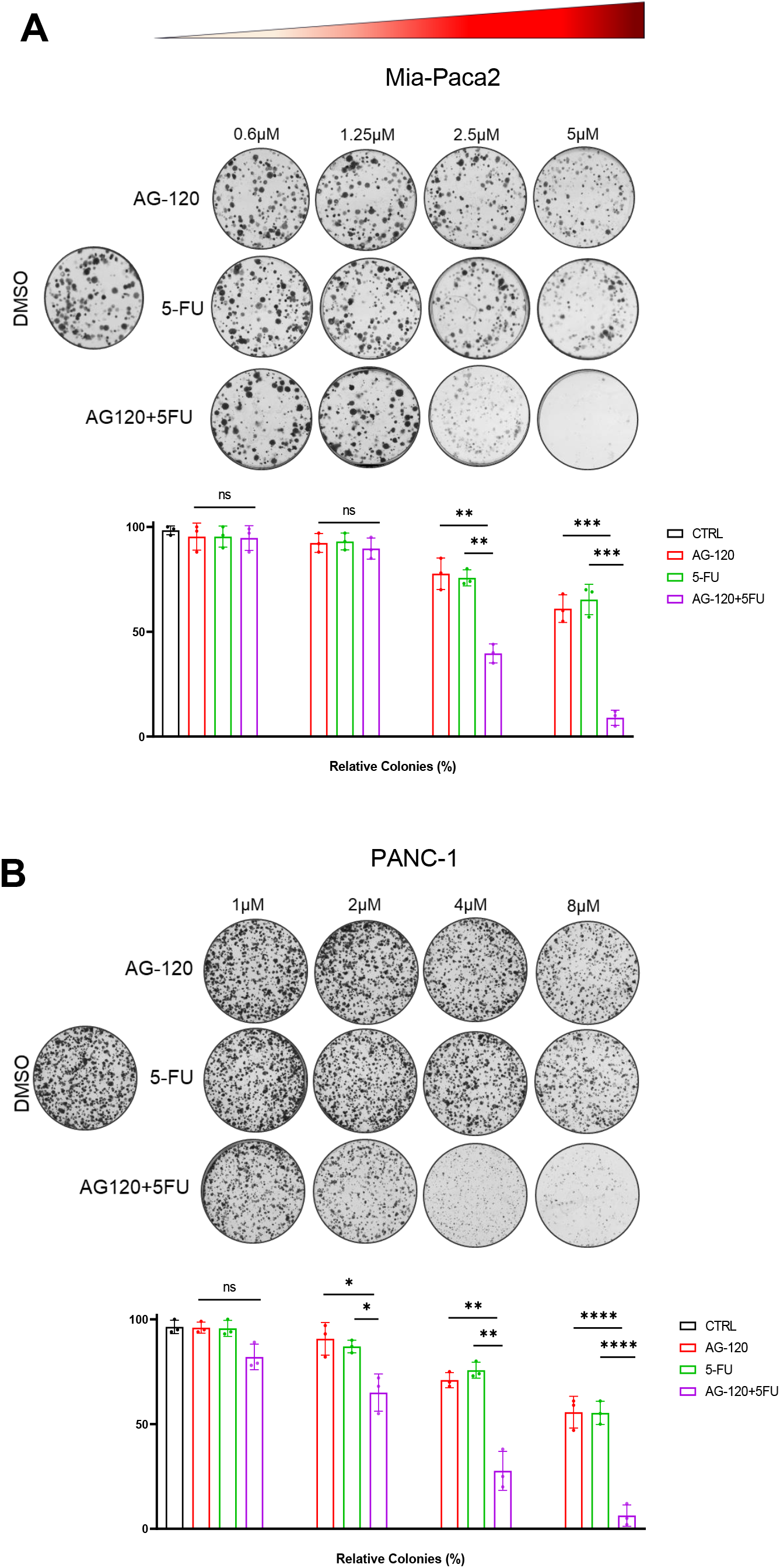
IDH1 inhibition decreases colony formation in combination with 5-FU in PC cells. Mia-Paca2 **A]** and PANC-1 **B]** were treated in increasing concertation of AG-120 and 5-FU (μM) alone or in combination, and the resulting colonies were later fixed and stained with 0.5% crystal violet solution. Bar graphs represent quantification of relative absorbance following extraction of crystal violet dye. Each data point represents the mean ± SEM of at least three independent experiments. N.S., nonsignificant; *, P < 0.05; **, P < 0.01; ***, P < 0.001; ****, P < 0.0001.

### Mitochondrial function is altered in PC cells treated with chemotherapy

Mitochondria play a central role in energy production and respiration (oxidative phosphorylation), and any alteration in mitochondrial cell metabolism typically results in downstream effects on cancer cell survival. Here we asked how cancer cell changes are associated with oxidative phosphorylation while treating cells with 5-FU and oxaliplatin allows cells to survive in the harsh microenvironment. The oxygen consumption rate (OCR) measured by the seahorse analyzer revealed increases in basal oxygen rate, ATP production and maximal mitochondrial respiration in PC cells treated with an IC_50_ dose of 5-FU and oxaliplatin under low glucose conditions **(Fig. 4A and B and Supplementary Fig. S2A)**. These data suggest that under nutrient deprived conditions, PC cells enhanced mitochondrial respiration when exposed to chemotherapy, suggesting this may be a pro-survival adaptation.

**Figure-4.**
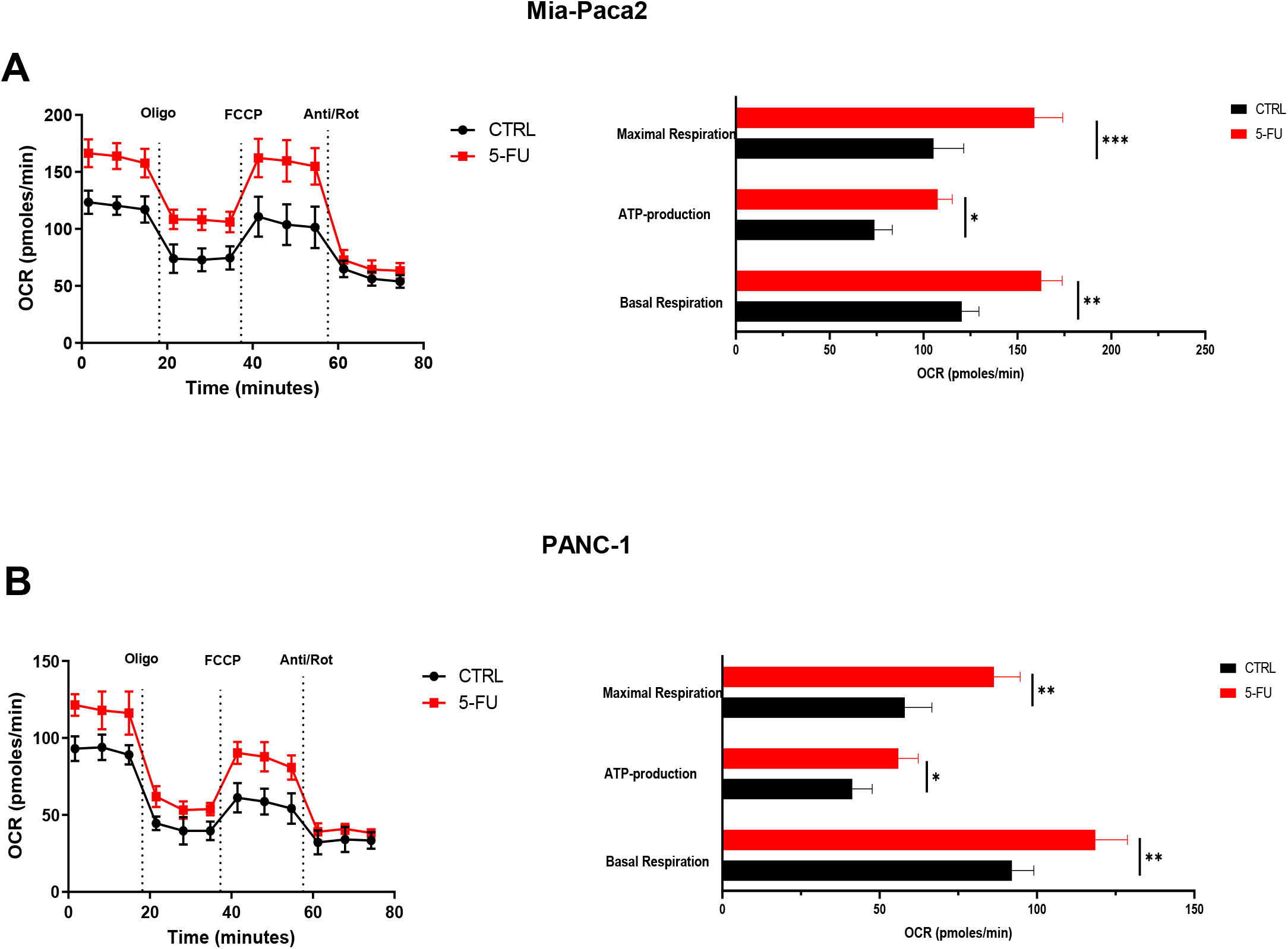
Mitochondrial function is altered in PC cells treated with chemotherapy. Mia-Paca2 **A]** PANC-1 **B]** Representative oxygen consumption rate (OCR) in PC cell lines cultured in 2.5 mM glucose treated with 5-FU for 36 hours. Treatment with mitochondrial inhibitors is indicated: oligomycin (Oligo), cyanide-p-trifluoromethoxyphenyl-hydrazone (FCCP), antimycin A and rotenone (Anti/Rot) and Basal mitochondrial respiration, ATP production, and maximal mitochondrial respiration. Each data point represents the mean ± SEM of three independent experiments. N.S., nonsignificant; *, P < 0.05; **, P < 0.01; ***, P < 0.001.

### IDH1 inhibition enhances 5-FU-induced cell apoptosis through ROS facilitated damage

Mitochondria are a source of endogenous ROS. As we previously observed increases in oxidative phosphorylation, we asked whether these increases were associated with ROS production. There was an increase in ROS when PC cells were treated with chemotherapy **(Fig. 5A and B)**. These data suggest that 5-FU drives ROS production and oxidative phosphorylation in PC cells.

**Figure-5.**
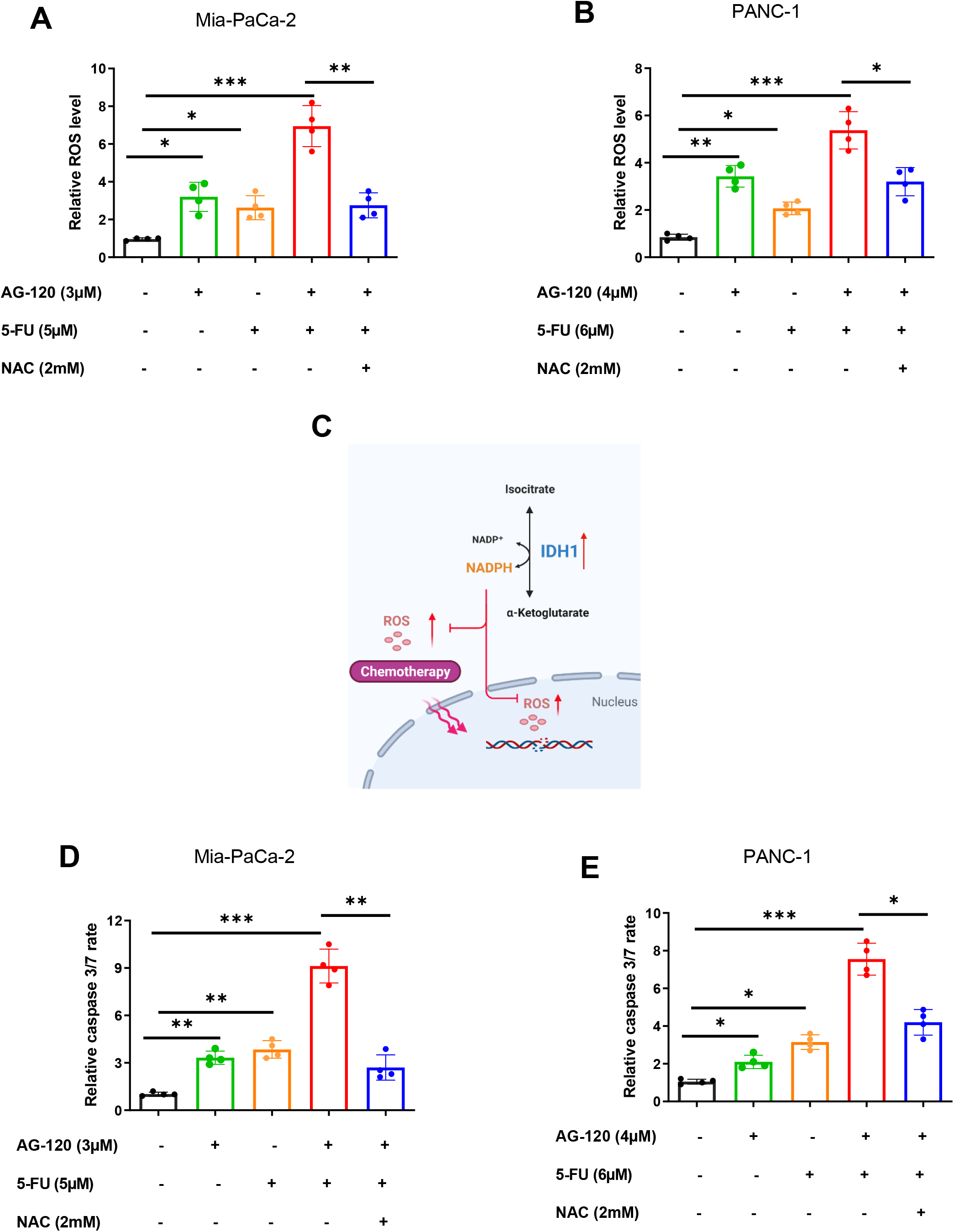
Combination of AG-120 and 5-FU induced apoptosis through ROS levels in PC cell lines. **A and B]** ROS levels were measured in Mia-Paca2 and PANC-1 cell lines by DCFDA after 48 hours of treatment with AG-120, 5-FU, AG-120 and 5-FU, or AG-120 and 5-FU with NAC rescue. **C]** Schematic of the IDH1 enzymatic reaction. Both products of wtIDH1 are vital for cancer cell survival under oxidative stress to support mitochondrial function and redox homeostasis. This represents how the combination of AG-120 and 5-FU induces apoptosis in PC cell lines. **D and E]** Caspase3/7 activity of control, AG-120, 5-FU, AG-120 and 5-FU, and AG-120 and 5-FU with NAC was measured in Mia-PaCa2 and PANC-1 cells after 48 hours. Each data point represents the mean ± SEM of at least three independent experiments. *, P < 0.05; **, P < 0.01; ***, P < 0.001; ****, P < 0.0001.

To further elucidate the underlying mechanism of synergy between IDH1 inhibition and 5-FU in PC cells, we assessed ROS levels in cells treated with 5-FU in combination with AG-120. As is shown in Figures 5A and B, 5-FU had an effect on ROS levels after 48 hours of incubation. When combined with AG-120, ROS levels increased by 3-fold in both PC cell lines. Importantly the effect of IDH1 inhibition and 5-FU combination was reversed by adding an antioxidant, N-acetylcysteine, a glutathione precursor (NAC; 2 mM) (**Fig. 5A and B)**. This could be a possible mechanism of synergistic effect between 5-FU and AG-120 IDH1, via the production of NADPH and αKG, rescues PC cells from chemotherapy damage **(Fig. E)**.

The ROS production overwhelms the cancer cells detoxification mechanism, ROS cytotoxicity damages DNA and cell membrane, resulting in cell death (33). Further, quantification of apoptosis (via Caspase3/7 activity) confirmed that IDH1 inhibition significantly increased the sensitivity to 5-FU after 48 hours of incubation. As with ROS levels, supplementation with NAC (2 mM) resulted in a decrease in apoptosis activity **(Fig. 5C and D).** These results propose that the accumulation of ROS contributes to the cytotoxic effect induced by AG-120 and 5-FU treatment, mediated through Caspase3/7 activation.

### IDH1 inhibition increases pancreatic cancer sensitivity to chemotherapies *in vivo*

The efficacy of AG-120 and 5-FU was explored *in vivo*. Nude mice bearing PDX (TM01212) were treated with vehicle, AG-120, 5-FU, or AG-120 and 5-FU. Mice receiving the combination of AG-120 and 5-FU had significantly smaller tumors as compared to all other treatment arms (**Fig. 6A-D**). Importantly, mice did not exhibit any signs of treatment-related toxicity, as body weights remained stable **(Fig. 6E).** Ki-67 immunolabeling of harvested tumors showed the proliferation rate was markedly decreased in mice receiving combination treatment **(Fig. 6F).** Furthermore, the combination of AG-120 and 5-FU resulted in more apoptosis by comparison with AG-120 or 5-FU, as was determined by cleaved caspase-3 (**Fig.6G**). Overall, the combination of AG-120 and 5-FU exhibited dramatic synergistic effects for pancreatic cancer treatment in PDX model.

**Figure-6.**
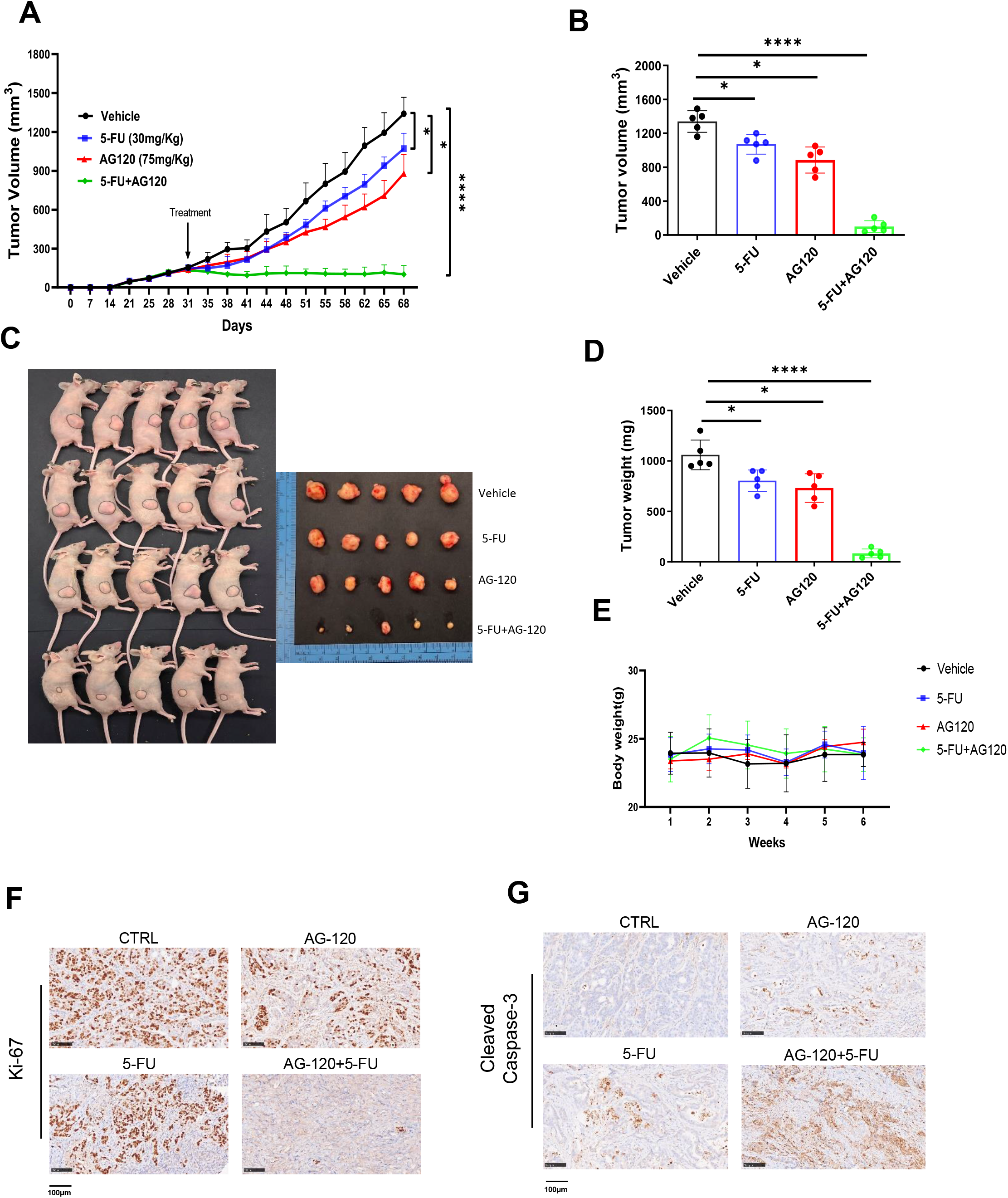
IDH1 inhibition increases PC sensitivity to 5-FU *in vivo.* **A]** Tumor growth curves of PDX (TM01212) in nude mice treated with vehicle, AG-120, 5-FU, or AG-120 and 5-FU for a total of 37 days. Treatment was started when tumors reached 100-120mm^3^ in size. Tumor volumes were calculated twice per week using calipers (n=5 per group). **B]** Average tumor volume at the end of the experiment (day 68) (n=5 tumors per group). **C]** Representative image of in vivo and excised PDX tumors at the end of the experiment. **D]** Average tumor weights (mg) of PDX tumors in each group (n=5 per group). **E]** Body weights of nude mice bearing PDX (n=5 per group). The paraffin-embedded tissue sections derived from nude mice bearing TM01212 PDX. **F]** Cell proliferation in PDX tumors was assessed by nuclear immunolabeling by Ki-67. Scale bar, 100 μm. **G]** Apoptosis was assessed in PDX tumors with labeled cleaved caspase-3. Scale bar, 100 μm. Each data point represents the mean ± SEM of at least three independent experiments. N.S., nonsignificant; *, P < 0.05; **, P < 0.01; ***, P < 0.001; ****, P < 0.0001.

Additionally, C57BL/6J mice bearing orthotopic pancreatic tumors received similar therapies. The first set of experiment mice were treated with both 5-FU and AG-120 and in a separate experiment mice were treated with both oxaliplatin and AG-120. In both sets of animal experiments there were significant improvements in median survival relative to monotherapies and vehicle (**Fig 7A and B**). Further to confirm our finding with FOLFIRINOX the standard care of treatment for pancreatic cancer patients. Nude mice bearing human pancreatic cancer Mia-paca2 cells were treated with vehicle, AG-120, FOLFIRINOX, or AG-120 and FOLFIRINOX. Mice treated with the combination of AG-120 and FOLFIRINOX had most effective as evidence observed in both tumor growth and tumor weight **(Fig. 7C and D)**. The combination of drugs was well tolerated as body weight remained stable during the entire treatment **(Fig. 7 E).**

**Figure-7.**
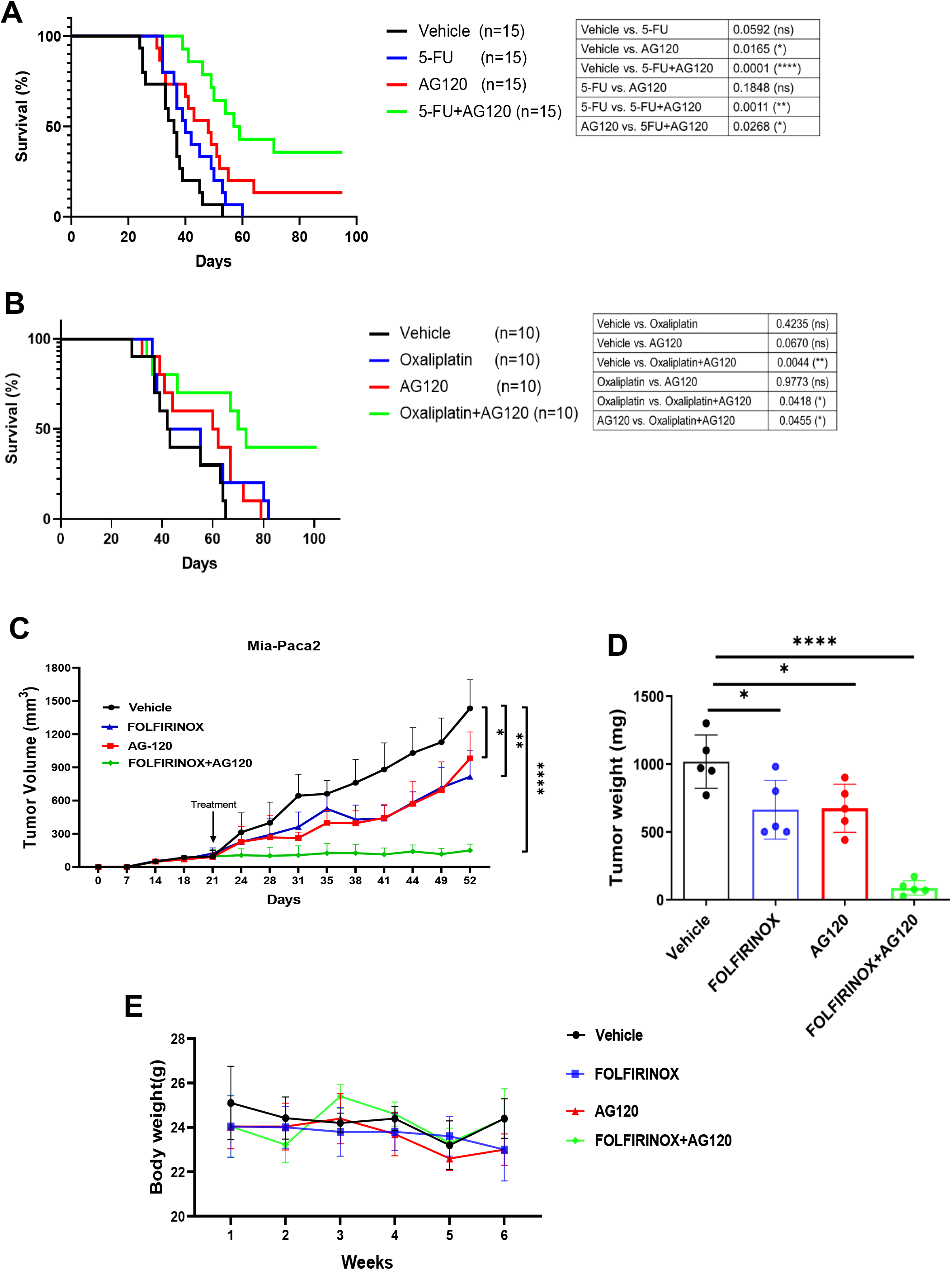
IDH1 inhibition increases PC sensitivity to chemotherapies *in vivo*. **A] and B]** Survival analysis of C57BL/6J mice bearing orthotopic pancreatic tumors, treated with (vehicle, AG-120, 5-FU, or AG-120 and 5-FU) and (vehicle, AG-120, oxaliplatin, or AG-120 and oxaliplatin). Median survival was analyzed using the Kaplan-Meier estimate and compared by the log-rank test. **C]** Tumor growth curves of Mia-Paca2 cells in nude mice treated with vehicle, AG-120, FOLFRIINOX, or AG-120 and FOLFIRINOX for a total of 32 days. Treatment was started when tumors reached 100-120mm^3^ in size. Tumor volumes were calculated twice per week using calipers (n=5 per group). **D]** Average tumor weights (mg) of tumors in each group (n=5 per group). **E]** Body weights of nude mice bearing (n=5 per group)

## Discussion

Tumor cells depend on metabolic alterations and other molecular strategies to neutralize oxidative stress and grow in extremely harsh metabolic microenvironments (34). We demonstrate that wild-type IDH1 up-regulation plays an important role in PC survival, and this enzyme represents a promising therapeutic target towards this end. Indeed, only a few studies highlight this enzyme as a convincing therapeutic target. The first published study by Metallo et al. observed that when cancer cells are cultured in hypoxic conditions, wtIDH1 activity increases, principally in the reductive direction: αKG derived from glutamine is converted to isocitrate, as an early step towards *de novo* lipogenesis. Importantly, suppression of this enzyme reduced cell growth (13). Later similar observations by Jiang et al. showed the importance of wtIDH1 activity in a reductive direction in tumor spheroid growth of different cancer models (11). Recent study by Calvert et al. demonstrated that the wtIDH1 reaction favors oxidative decarboxylation to support glioblastoma tumor growth by producing NADPH as the main antioxidant defense mechanism to control ROS. Suppression of this wtIDH1 resulted in the augmentation of oxidative stress and tumor growth inhibition (35).

Our recent study validated the importance of wtIDH1 in PC and established several key principles in that work. Cancer cells depend on wtIDH1 for generation of NADPH and αKG (produced by the oxidative decarboxylation of isocitrate) for adaptive survival under nutrient-limiting conditions (14–16,36). These two products play a role in antioxidant defense mechanisms (NADPH) and mitochondrial function (αKG) to support cancer cells under metabolic stress. We show that AG-120, a small molecule originally developed to target mutant IDH1, is actually a potent wtIDH1 inhibitor in PC tumors due to low Mg^2+^ levels, which permit stronger binding of the compound within the allosteric site of wtIDH1, and low nutrient concentrations which increases cancer cell dependency on this enzyme (14,36). These common conditions in the tumor microenvironment render cancer cells vulnerable to IDH1 inhibition.

Current cytotoxic chemotherapies are relatively ineffective against PC. This chemoresistance is likely due to rapid upregulation of compensatory pathways (37). Here, for the first time, we demonstrate that cellular NADPH and αKG, through the action of wtIDH1, are important for the efficacy of chemotherapy against PC. 5-FU, a common component of modern multi-agent regimens for patients with PC, is a uracil analog. It functions by intracellular conversion to several active metabolites that disrupt RNA synthesis and thymidylate synthase, causing cytotoxicity (38). We investigated the role of wtIDH1 in drug resistance and observed that treatment of PC cells with 5-FU resulted in increases in ROS and apoptosis, with an induction of wtIDH1. Our data show that intracellular ROS is important for 5-FU cytotoxicity. Increases in ROS usually results in a compensatory increase in HuR and NRF-2. These compensations initiate an antioxidant defense mechanism to prevent cellular injury by ROS, ultimately resulting in chemoresistance (15,39). The excessive level of ROS can exhaust the antioxidant capacity and promote apoptosis (40). Combining the AG-120 and 5-FU resulted in increased apoptosis through a high level of ROS **(Fig. 5)**. Herein, we systematically show that wtIDH1 inhibition with AG-120 dramatically sensitizes PC to treatment with chemotherapy. AG-120 has been shown to quite safe with minimal side effects, suggesting this therapeutic combination could be used for patients with PC.

In conclusion, our results highlight the effectiveness of AG-120 combined with chemotherapy, which increased ROS generation and PC cell apoptosis. Compensatory upregulation of wtIDH1 provides a novel explanation of why chemotherapy may be ineffective as a monotherapy in patients with PC. Further understanding the role of wtIDH1 in PC may allow it to be explored as a prognostic biomarker for predicting response to based chemotherapy and aid in the development of novel therapeutic strategies.

## Abbreviations

PDAC: Pancreatic ductal adenocarcinoma
IDH1: Isocitrate dehydrogenase 1
5-FU: 5-Fluorouracil
αKG: alpha-ketoglutarate
TCGA: The Cancer Genome Atlas
TCA: Tricarboxylic acid cycle
ROS: Reactive oxygen species

## Authors’ Contributions

**M. Zarei:** Investigation, data curation methodology, writing-original draft, writing–review and editing. **O. Hajihassani:** Investigation, methodology. **J. J. Hue:** Investigation, methodology. **H. J. Graor:** Investigation, methodology. **L. D. Rothermel:** funding acquisition, Investigation, methodology. **J.M. Winter:** Conceptualization, funding acquisition, supervision, administration, writing and editing.

